# A single polymorphic residue in humans underlies species-specific restriction of HSV-1 by the antiviral protein MxB

**DOI:** 10.1101/2023.05.30.542951

**Authors:** Avraham Bayer, Stephanie J. Child, Harmit S. Malik, Adam P. Geballe

## Abstract

Myxovirus resistance proteins (MxA and MxB) are interferon-induced proteins that exert antiviral activity against a diverse range of RNA and DNA viruses. In primates, MxA has been shown to inhibit myxoviruses, bunyaviruses, and hepatitis B virus, whereas MxB restricts retroviruses and herpesviruses. As a result of their conflicts with viruses, both genes have been undergoing diversifying selection during primate evolution. Here, we investigate how MxB evolution in primates has affected its restriction of herpesviruses. In contrast to human MxB, we find that most primate orthologs, including the closely related chimpanzee MxB, do not inhibit HSV-1 replication. However, all primate MxB orthologs tested restrict human cytomegalovirus. Through the generation of human and chimpanzee MxB chimeras we show that a single residue, M83, is the key determinant of restriction of HSV-1 replication. Humans are the only primate species known to encode a methionine at this position, whereas most other primate species encode a lysine. Residue 83 is also the most polymorphic residue in MxB in human populations, with M83 being the most common variant. However, ∼2.5% of human MxB alleles encode a threonine at this position, which does not restrict HSV-1. Thus, a single amino acid variant in MxB, which has recently risen to high frequency in humans, has endowed humans with HSV-1 antiviral activity.

**Importance:** Herpesviruses present a major global disease burden. Understanding the host cell mechanisms that block viral infections as well as how viruses can evolve to counteract these host defenses is critically important for understanding viral disease pathogenesis, and for developing therapeutic tools aimed at treating or preventing viral infections. Additionally, understanding how these host and viral mechanisms adapt to counter one another can aid in identifying the risks of, and barriers to, cross-species transmission events. As highlighted by the recent SARS-CoV-2 pandemic, episodic transmission events can have severe consequences for human health. This study reveals that the major human variant of the antiviral protein MxB inhibits the human pathogen HSV-1, whereas human minor variants and orthologous MxB genes from even closely related primates do not. Thus, in contrast to the many antagonistic virus-host interactions in which the virus is successful in thwarting the defense systems of their native hosts, in this case the human gene appears to be at least temporarily winning at this interface of the primate-herpesviral evolutionary arms race. Our findings further show that a polymorphism at amino acid 83 in a small fraction of the human population is sufficient to abrogate MxB’s ability to inhibit HSV-1, which could have important implications for human susceptibility to HSV-1 pathogenesis.

## Introduction

Upon exposure to interferon, cells generate a robust antiviral response composed of hundreds of different interferon stimulated genes (ISGs) capable of restricting different stages of the viral lifecycle [1]. Some ISGs differentially target specific viruses, while others exhibit a broader range of restriction. This antiviral landscape has been shaped by interactions with myriad viral pathogens over evolutionary time. Viral adaptations that enable evasion of the cellular defenses exert pressure on the host genes to evolve and refine their antiviral potency. The ongoing reciprocal evolution of the host and virus is consistent with the Red queen genetic conflict hypothesis [2, 3] and leads to an increased rate of nonsynonymous (amino acid altering) compared to synonymous (amino acid preserving) nucleotide substitutions among host restriction factor homologs in related host species.

One such rapidly evolving interferon-induced antiviral factor is myxovirus resistance protein B (MxB; also known as Mx2) [4–6]). MxB is one of two dynamin-like GTPase antiviral ISG proteins. Previous studies showed that primate MxB is an important restriction factor of HIV-1 [7, 8]. However, the rapidly evolving residues in MxB do not correspond to those that were identified as important for lentiviral restriction [6]. Thus, the recent finding that MxB is also a pan-herpesvirus restriction factor [9–11] raised the possibility that the rapid evolution of MxB might instead result from pressure to evade herpesviruses. To test this possibility, Schilling et al. tested a few substitutions at positively-selected residues but found none that impacted the ability of human MxB to inhibit herpes simplex virus (HSV-1) replication [10]. However, it remained possible that the primate MxB orthologs differ in their ability to restrict HSV-1 due to variants that were not tested or to epistatic interactions between multiple residues that differ between the orthologs. Moreover, even if HSV-1 and related herpesviruses did not drive the evolution of MxB, they would nevertheless have been under selection to evade MxB restriction.

Here, we investigated whether the evolution of MxB has affected its potential for inhibiting herpesviruses. We compared the ability of human, chimpanzee, and representative Old World and New World monkey MxB orthologs to restrict herpesvirus replication. We found that all of the tested primate MxB orthologs could inhibit cytomegalovirus (HCMV), but that only human MxB restricted HSV-1. By generating and testing human-chimp MxB chimeras we were able to identify a single residue, M83, as the critical determinant of HSV-1 sensitivity. While most human MxB alleles encode a methionine at residue 83, codon 83 is a lysine in chimps and most other primates. In fact, a K83M substitution in chimp MxB was sufficient to confer HSV-1 restriction. We also found that although M83 is unique and highly prevalent in human MxB alleles, a minor fraction of human MxB alleles encode a threonine at this position, which corresponds to a loss of HSV-1 restriction. Our work identifies a single residue change that confers HSV-1 restriction in human MxB, and a loss-of-function minor allele polymorphism that could affect HSV-1 disease pathogenesis.

## Results

### MxB expression is necessary for HSV-1 inhibition in fibroblasts

Previous studies showed that HSV-1 titers increased upon MxB knockdown in interferon (IFN)-treated cells [9, 10]. We recapitulated these results in IMR90 cells by transducing them with a lentiviral vector expressing Cas9 and an MxB-targeting guide RNA or a non-targeting control (NTC). We either pretreated these cells with IFN for 24h or left them untreated and assessed MxB expression by immunoblotting (Fig. 1A). These results confirmed a lack of MxB expression in the KO cells.

**Fig 1.**
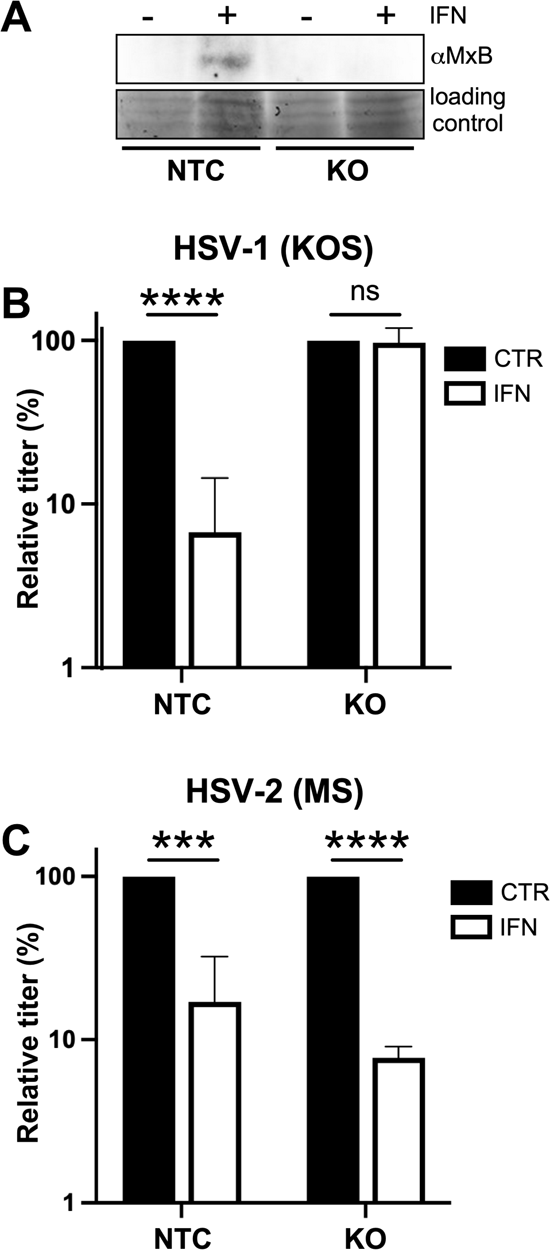
MxB expression is necessary for HSV-1 inhibition in fibroblasts. (A) MxB knock out IMR90 fibroblasts (KO) or non-targeting control (NTC) cells were left untreated or treated with 1000 units/ml IFN β for 24h. Cell lysates were then collected, and MxB expression was analyzed by immunoblot assay. (B) The cells described in (A) were left untreated or treated with IFN β, infected with HSV-1 (strain KOS; MOI = 0.1) for 48h, then the amount of virus present in the medium was determined by plaque assay. (C) Cells treated as in (B) were infected with HSV-2 (strain MS; MOI = 0.1) for 48h and the virus in the medium was titered as in (B). Titers were normalized to those in the untreated controls. Statistical significance for replication in control vs KO cells was determined using a Student’s T test (ns P>0.05; *** = P<0.001; **** = P<0.0001).

We next treated the KO and NTC cells +/- IFN and then infected them with HSV-1 (strain KOS) at an MOI of 0.1. As expected, IFN treatment reduced HSV-1 replication in the control NTC cells, yet we observed a >10-fold restoration of viral titers in the MxB KO cells (Fig. 1B). These results suggest that MxB is a major component of the ISG response to HSV-1 in IMR90 cells. When the same cells were infected with HSV-2 (strain MS, MOI = 0.1), equivalent levels of restriction were observed in the control and KO cells. These results confirm that the IFN system is functional in IMR90 cells, and reveal that, unlike with HSV-1, MxB is not necessary for restriction of HSV-2 replication (Fig. 1C). These data demonstrate that MxB is necessary for the restriction of HSV-1 but not HSV-2 in interferon-treated IMR90 fibroblasts.

### Restriction of HSV-1 by MxB is species-specific

*MxB* has been evolving under diversifying selection throughout primate evolution (and likely in all mammals), which suggests it has been engaged in arms races with a changing repertoire of viruses. We tested whether one consequence of this lineage-specific engagement with different viruses may have led to differences in primate MxB abilities to inhibit herpesviruses. We transduced our MxB-KO IMR90 cell line with lentiviruses containing either a doxycycline (dox)-inducible human *MxB* or *MxB* orthologs from representative simian primates: *Pan troglodytes* (chimpanzee, Hominin), *Chlorocebus tantalus* (African Green monkey, Old World Monkeys), and *Aotus trivirgatus* (owl monkey, New World monkeys). None of these *MxB* orthologs can be targeted by the guide RNA sequence used to engineer the KO cells since they each contain synonymous mutations at the target site.

We confirmed that doxycycline treatment induced expression of the *MxB* orthologs (Fig. 2A) although expression levels varied among the orthologs. To test whether HSV-1 was sensitive to the different MxB orthologs, we infected dox-treated or untreated cells with HSV-1 (strain KOS, MOI of 0.1), and titered progeny virus harvested at 48 hpi. Surprisingly, we found that only the human MxB ortholog restricted HSV-1 (Fig. 2B). We confirmed that the restriction was not strain-specific, as HSV-1 strain 17 demonstrated a similar phenotype to KOS (Fig. 2C). We also infected these cells with human cytomegalovirus (HCMV strain AD169, MOI = 0.1) and quantified progeny production at 6 days pi (Fig. 2D). In contrast to HSV-1, we observed that all MxB orthologs restricted HCMV, despite low levels of expression induction in some cases. Thus, *MxB* restriction of HSV-1, but not HCMV, appears to be species-specific under these conditions.

**Fig 2.**
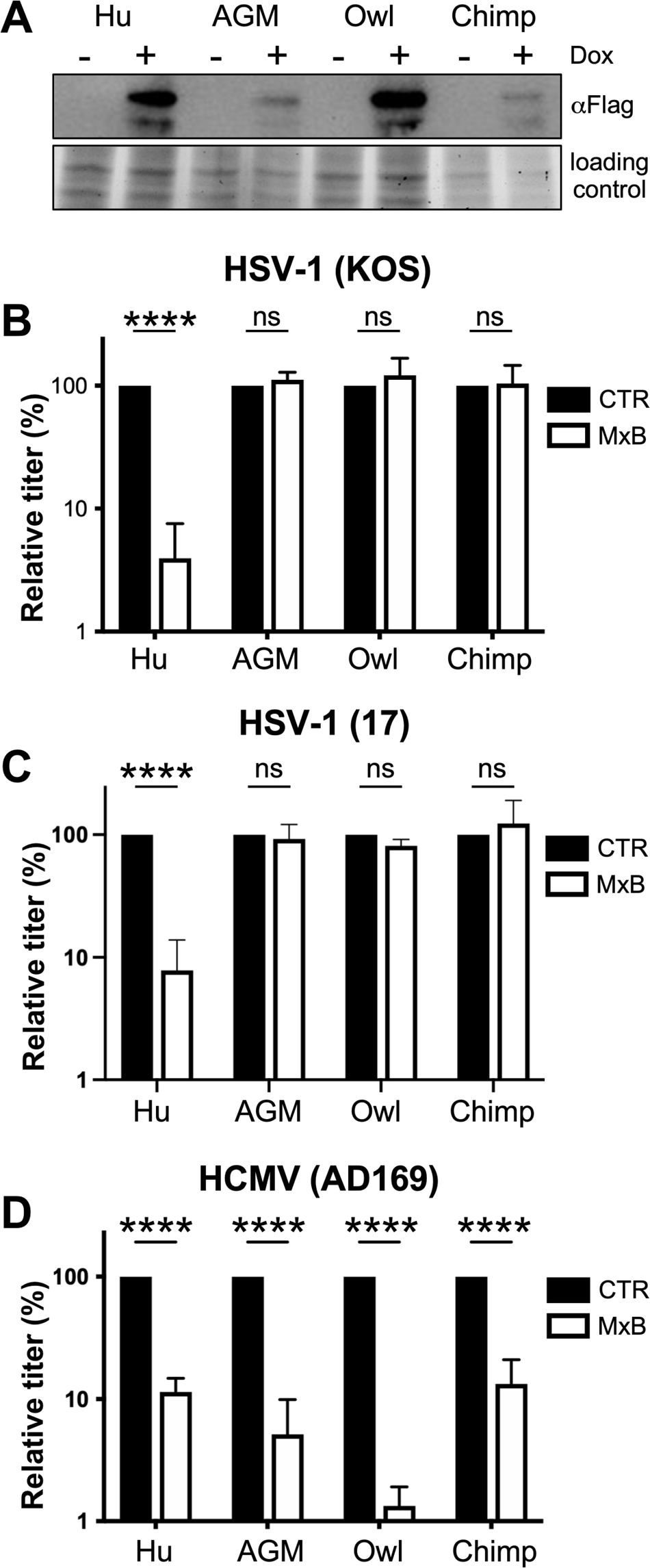
Restriction of HSV-1 by MxB is species-specific. (A) Immunoblot analysis of cell lysates from IMR90s expressing the MxB orthologs (human, chimpanzee, African green monkey, or owl monkey) that had been untreated or treated with dox (1 µg/ml) for 24h to determine MxB expression levels. The ortholog-expressing IMR90s treated as in (A) were infected with HSV-1 strain KOS (B) or HSV-1 strain 17 (C) for 48h (MOI = 0.1). Virus present in the medium was then titered, and the titers normalized to the untreated controls. (D) The cells were treated as in (B), then infected with HCMV (strain AD169; MOI = 0.1) for 6 days, following which the virus in the medium was titered and the titers normalized as above (ns P>0.05; ** = P ⊔ 0.01; **** = P<0.0001).

### Species specificity of nonhuman primate MxB is not due to poor expression or mis-localization

Although the observed differences in expression of the MxB orthologs in the IMR90 cells did not eliminate their ability to inhibit HCMV (Fig. 2A,D), it remained possible that this variable expression might contribute to the deviation in their impact on HSV-1. To eliminate integration site variation, a potential contributor to variability in transgene expression, we introduced the *MxB* orthologs into the landing pad using Cre-Lox recombination in U2OS cells [12]. Expression of the *MxB* orthologs was less variable using this strategy (Fig. 3A). However, infection experiments in U2OS cells confirmed our previous finding of species-specific restriction of HSV-1 by *MxB* in IMR90 cells: only cells expressing human *MxB*, but none of the other orthologs, restricted HSV-1 replication (Fig. 3B).

**Fig. 3.**
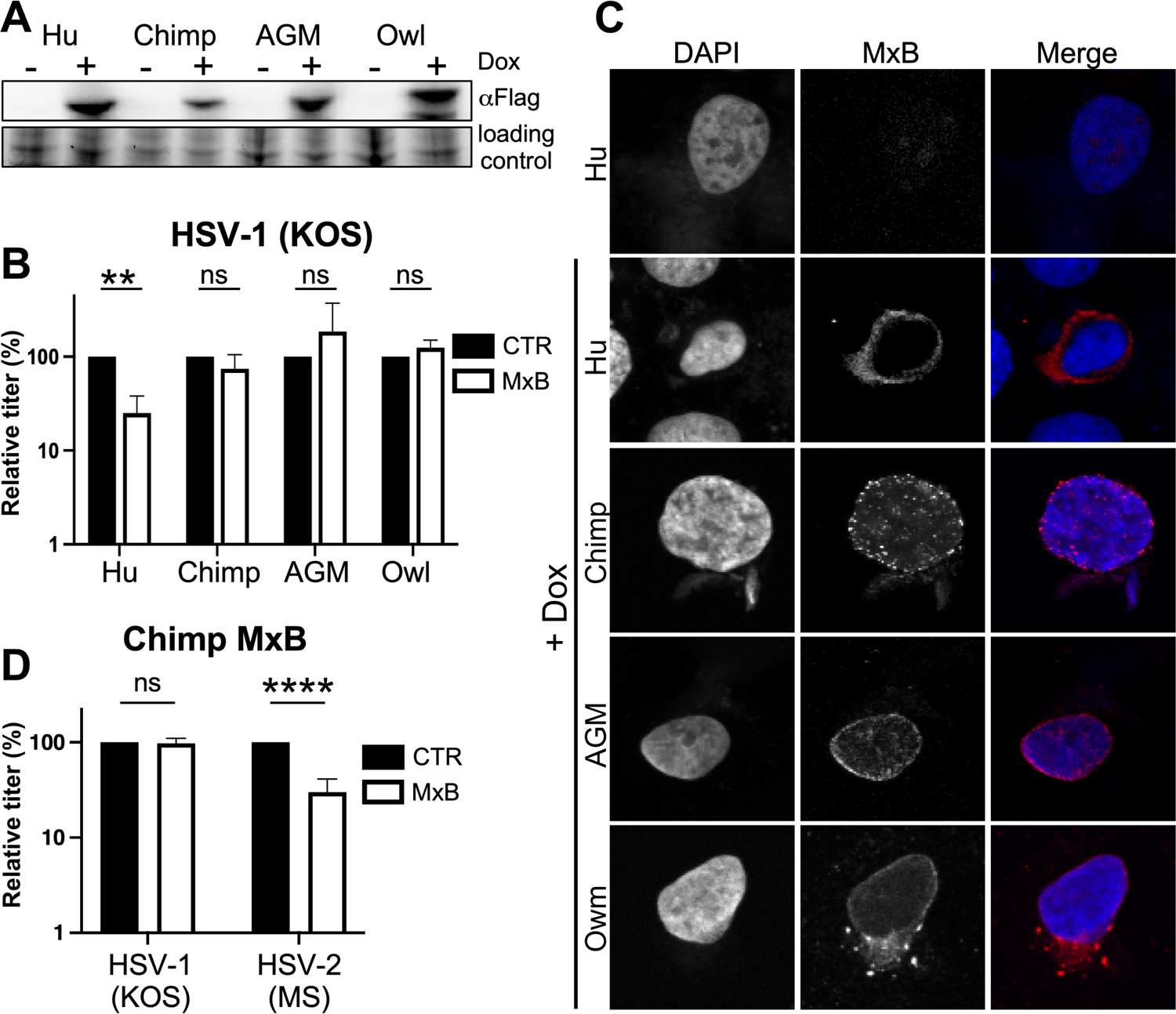
Species-specificity of the nonhuman primate MxB orthologs is not due to poor expression or mis-localization. (A) Immunoblot analysis of cell lysates from U2OS cells expressing the different MxB orthologs (untreated or treated with dox (1 µg/ml) for 24h to induce MxB expression) to determine the expression level of each ortholog. (B) These U2OS cells, either untreated (CTR) or treated with dox (MxB) were infected with HSV-1 (strain KOS; MOI = 0.1) for 48h, and titers quantified and normalized to the untreated controls as described above. (C) The MxB-expressing cells were fixed with 4% PFA and stained to detect flag-tagged MxB (red) or DNA (Blue), then imaged by confocal microscopy. (D) The chimp MxB-expressing U2OS cells (treated or untreated) were infected with either HSV-1 (KOS; MOI = 0.1) or HSV-2 (MS; MOI = 0.1), and supernatant virus collected and quantified as in the preceding figures (ns P>0.05; ** = P⊔ 0.01; **** = P<0.0001).

Prior studies have demonstrated that MxB localizes to the cytoplasmic face of the nuclear membrane, and associates with nuclear pore complex components, such as NUP358 [13]. Indeed, this subcellular localization of MxB, dependent on the nuclear localization signal present in the N-terminus, has been documented as being critical for restriction of lentiviruses and herpesviruses [10, 14, 15]. We therefore evaluated whether the nonhuman MxB orthologs also localized to the nuclear membrane, or if their inability to counteract HSV-1 could be attributed to mislocalization. We treated the MxB variant expressing U2OS cells with dox (24h) to induce MxB expression, fixed the cells, and stained them with anti-flag antibody (Materials and Methods). Confocal microscopy of these samples revealed predominantly perinuclear expression pattern for all MxB orthologs (Fig. 3C). These results are consistent with previous reports describing Old World monkey MxB localization [5] and suggest that the observed species-specificity of HSV-1 restriction is likely not due to variation in expression or localization. Rather, another feature of the amino acid variation among MxB must account for their different abilities to restrict HSV-1 replication.

### Chimpanzee MxB can restrict HSV-2 but not HSV-1

To further explore the anti-herpesviral spectrum of the hominoid MxB orthologs, we tested whether chimp MxB could restrict HSV-2. Humans are the only primate species to be infected with two different simplex viruses; it is believed that HSV-2 evolved following the transfer of chimp simplex virus into humans ∼1.6 million years ago [16, 17]. The MxB-expressing U2OS cells were treated +/- dox for 24h, then infected with either HSV-1 (strain KOS) or HSV-2 (strain MS) at an MOI of 0.1. Supernatant virus was collected at 48h post-infection, and viral replication was measured by titering. As before, we observed that chimp MxB did not restrict HSV-1. However, chimp MxB was able to restrict HSV-2 replication by approximately 5-fold (Fig. 3D). Thus, both human and chimp MxB inhibit HCMV and HSV-2 [9, 10], whereas only human MxB is capable of inhibiting HSV-1, further suggesting that sequence variation among these MxB orthologs confers species-specific differences in their antiviral spectra.

### Codon 83 is a critical determinant of HSV-1 restriction

Human and chimp MxB share 98% sequence identity, differing by 17 nucleotides, only 11 of which encode nonsynonymous substitutions. Thus, we were particularly intrigued by the observation that human, but not chimp, MxB is able to restrict HSV-1 (Fig. 2B and C; Fig. 3 B and D). To map the determinants responsible for the differential effect of human and chimp MxB on HSV-1, we first made six chimeric human-chimp constructs (Fig. 4A), introduced them into the U2OS landing pad cells, then tested them for restriction of HSV-1. All chimeras containing the N-terminal portion of human MxB were capable of restricting HSV-1 (constructs Fig. 4B, chimeras B, C, and E) whereas those containing the chimp N-terminal region did not (Fig. 4B, chimeras A, D, and F).

**Fig. 4.**
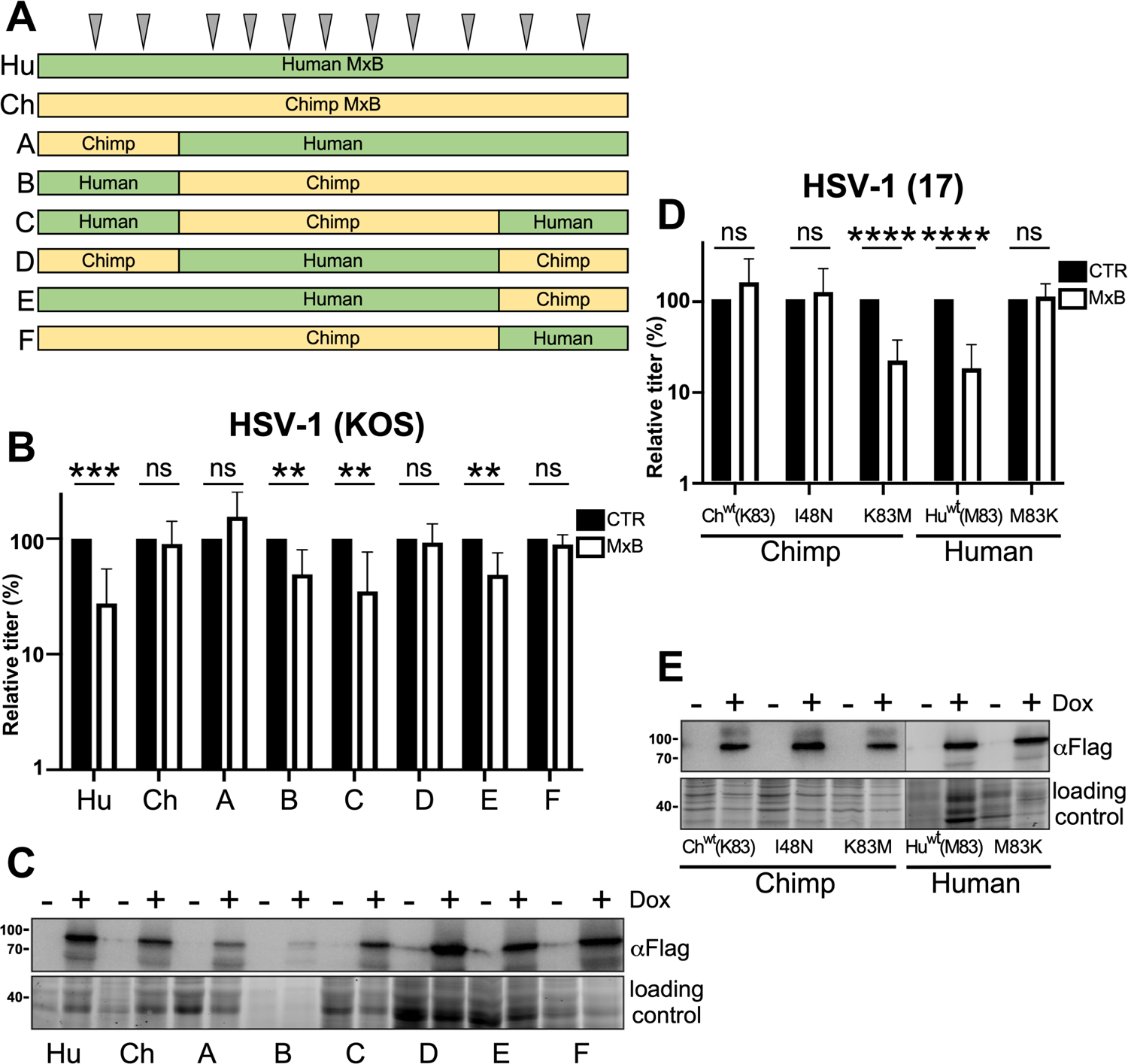
Codon 83 is a critical determinant of HSV-1 restriction. (A) Schematic of human MxB, chimp MxB, and the human-chimp chimeras. Grey arrows indicate nonsynonymous substitutions between human and chimp MxB. (B) Titers of HSV-1 (KOS; MOI = 0.1) present in the medium at 48 h after infection of U2OS cells expressing human, chimp, or each of the chimeric MxBs as described in previous figures (ns P>0.05; ** = P< 0.01; *** = P<0.001). (C) Immunoblot analysis to determine the expression levels of each of the MxBs shown in (A) in the presence (+) or absence (-) of dox. (D) Replication of HSV-1 (17; MOI = 0.1) was measured at 48 h post-infection in cells containing wild type chimp or human MxB, point mutants I48N and K83M in the chimp gene, or point mutant M83K in the human background following a 24h incubation with (MxB) or without (CTR) dox. Titering and normalization were performed as described in previous figures (ns P>0.05; **** = P<0.0001). (E) Immunoblot analysis to demonstrate expression levels of the MxBs described in (D).

Although different MxB chimeras varied in their expression (Fig. 4C), expression variation did not correlate with HSV-1 restriction. For example, chimeras C and E restricted HSV-1 replication despite being expressed poorly relative to chimera D, which could not restrict HSV-1. Thus, differences in the N-terminal domain of MxB account for the differences in the ability of the human and chimp genes to block HSV-1 replication. This N-terminal portion of MxB contains only two nonsynonymous substitutions between the human and chimp alleles at codons 48 and 83. We therefore engineered point mutations to test whether single human MxB amino acids would enable chimp MxB to inhibit HSV-1. Introduction of the human variant at position 48 (asparagine) into chimp MxB did not endow it with the ability to restrict HSV-1 (Fig. 4D and 4E). However, replacing lysine at residue 83 in chimp MxB with methionine from human MxB, was sufficient to enable chimp MxB to inhibit HSV-1 with an efficacy comparable to that of human MxB. Conversely, human MxB lost the ability to restrict HSV-1 upon replacing methionine 83 in human MxB with lysine (from chimp MxB) (Fig. 4D). These results suggest that a methionine at position 83 is necessary and sufficient to enable inhibition of HSV-1 by hominoid MxBs.

### Codon 83 variation does not affect protein stability

We wondered whether the inability of lysine83 variants of MxB may be related to virally-induced degradation by HSV-1, as has been reported for other HSV-1 restriction factors [18]. However, upon comparing the levels of human and chimp MxB in mock or HSV-1 infected cells, we observed that infection did not affect expression or either MxB protein, indicating that lysine 83 does not sensitize the ortholog to degradation (Fig. 5).

**Fig. 5.**
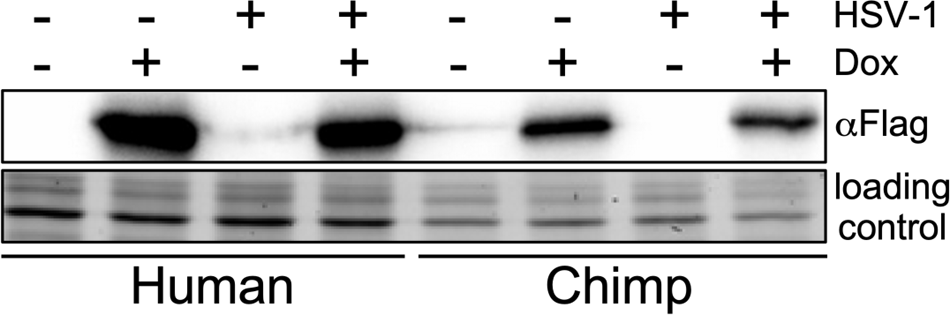
Codon 83 variation does not affect protein stability. (A) Measurement of MxB expression levels from human or chimp MxB-expressing cells that were untreated or pre-treated with dox, then either mock-infected or infected with HSV-1 (MOI = 5) for 24 h.

### Human-specific evolution of MxB variants

Our results indicate that a single human-specific non-synonymous variant that converted an ancestral lysine codon to a methionine codon is responsible for gain of HSV-1 restriction. We traced the evolutionary origins of this variant prior to the separation of human, Neanderthal, and Denisovan lineages; all of them encode a methionine at this residue 83.

However, this position is polymorphic within extant humans, with a M83T (methionine -> threonine) variant present at a minor allele frequency of 2.5% in all humans, but at an elevated allele frequency of ∼18% in African and African American populations (21-41377154-T-C variant, gnomAD browser [19, 20]). We tested whether this non-synonymous polymorphism affected MxB cytolocation or restriction of HSV-1. We found that the T83 variant was displayed a similar perinuclear localization (Fig. 6A) and was expressed to a similar level as wildtype MxB (Fig. 6B), and was able to restrict HSV-2 (Fig. 6C). However, the T83 variant was unable to restrict HSV-1 infection (Fig. 6D). These data confirm that although a recently arising methionine residue at position 83 of full-length MxB is critical for the anti-HSV-1 activity of human MxB, a significant fraction of humans have lost this restriction.

**Fig. 6.**
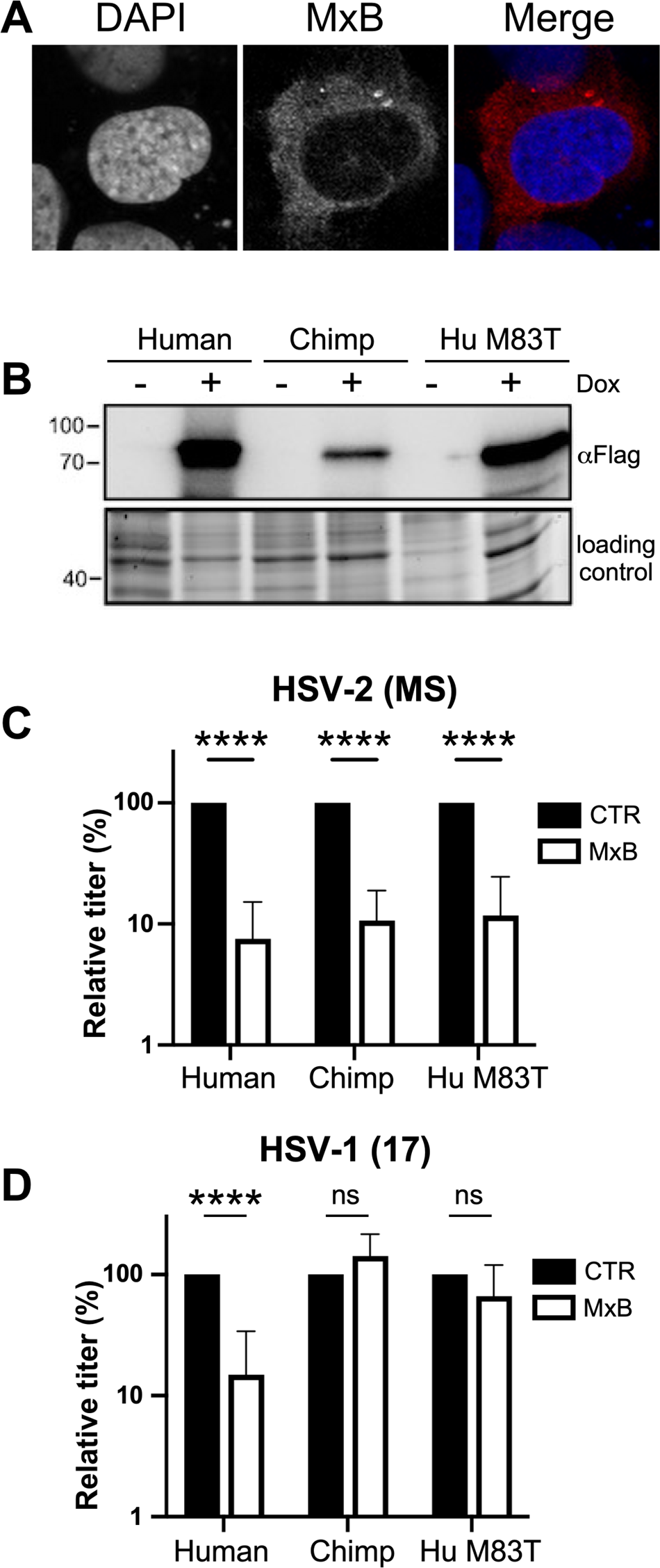
Human-specific evolution of MxB variants. (A) U2OS cells expressing the human M83T MxB variant were fixed with 4% PFA and stained to detect flag-tagged MxB (red) or DNA (Blue), then imaged by confocal microscopy. (B) Immunoblot demonstrating the expression levels of U2OS cells expressing the human, chimp, or human MxB M83T point mutant +/- dox. The untreated or dox-treated cells described in (B) were infected with (C) HSV-2 (MS; MOI = 0.1) or (D) HSV-1 (strain 17; MOI = 0.1) for 48h, following which titers were quantified and normalized as before (ns P>0.05; **** = P<0.0001).

## Discussion

Our work extends previous findings implicating the important role of MxB as a herpesvirus restriction factor in primates [9–11]. Our knockout experiments further reveal that MxB is the primary or only ISG affecting HSV-1 in IMR90 cells (Fig. 1), similar to previous findings in T98G fibroblast-like cells [10]. However, the extent to which herpesvirus replication is affected following interferon treatment differs between these previous reports and our findings. We attribute these differences to the variable expression of other ISGs between different cell lines. Regardless, the magnitude of MxB’s role in inhibiting HSV-1 (based on ∼10-fold restoration of HSV-1 titers in MxB KO cells) is similar to previous reports, supporting the conclusion that MxB is a potent inhibitor of HSV-1.

MxB is among the set of broadly-acting ISGs, which target multiple viruses [21]. Previous computational analyses revealed that 6 residues in MxB have undergone recurrent positive selection during simian primate evolution, presumably in response to one or more viral infections [5]. However, these sites did not overlap with the sites that have been demonstrated to be critical for HIV-1 restriction, suggesting that MxB rapid evolution was likely driven by pathogens other than lentiviruses. In response to the finding that MxB can inhibit herpesviruses [9–11], Schilling *et al.,* tested a few point mutations at rapidly evolving residues but found that they did not alter the restriction of HSV-1 by human MxB suggesting that HSV-1 is not affected by evolution at those sites [10].

Since recurrent positive selection requires multiple, independent episodes of adaptation at the same interaction interface, this may not represent all modes of adaptive evolution driven by host-virus arms races. Therefore, we undertook a more agnostic approach to test whether there is any variability in herpesviral restriction by primate MxB orthologs. We found that representative MxB orthologs from non-human primates (chimpanzee, African green monkey, owl monkey) were all able to restrict HCMV despite being variable at rapidly evolving residues. However, only human MxB was able to restrict HSV-1. This unique HSV-1 restriction ability of human MxB could be attributed to residue 83, which has not undergone recurrent positive selection [4–6]. Thus, our data do not support a role for herpesviruses having driven recurrent positive selection of primate MxB HSV-1.

However, selection for HSV-1 could have driven the single K83M change in MxB. A single, lineage-specific, episode of adaptation would not lead to a signature of recurrent positive selection. Moreover, since this change occurred in the common ancestor of humans, Neanderthals, and Denisovans, there is not enough standing variation to test whether this change was driven by an adaptive sweep at this locus. Nevertheless, the methionine at residue 83 is not observed in any other primate MxB, which are conserved for lysine instead. Our finding that a single residue is critical for species-specific antagonism by MxB is consistent with past work reporting that very few residues can confer species-specific lentiviral restriction by MxB. For example, human MxB but not African green monkey MxB restricts clade O HIV-1, with residues 37 and 39 identified as the determinants of these phenotypic differences [5]. Similarly, residue 37 accounts for the restriction of an HIV-1 capsid mutant by Rhesus macaque but not African green monkey MxB.

Our data confirm previous reports that human MxB restricts HSV-1, HSV-2, and HCMV. However, we find that HSV-1 restriction is unique to human MxB among primate orthologs. For example, while the chimp ortholog restricts HCMV and HSV-2, it does not restrict HSV-1. Humans are the only primate known to be infected with 2 herpes simplex viruses. HSV-2 crossed from the chimpanzee ancestral lineage into the human ancestral lineage about 1.6 million years ago, and is more closely related to extant chimpanzee simplex virus than to HSV-1 [16]. Therefore, chimp MxB may have been better adapted to restricting HSV-2 because of its co-evolution with chimpanzee simplex virus, but not with HSV-1. Nevertheless, it is intriguing that the apparent selection of M83 in the human ancestral MxB resulted in it acquiring the ability to inhibit HSV-1 without affecting its ability to counteract at least two other herpesviruses (HCMV and HSV-2).

What is the mechanism underlying the different HSV-1 restriction abilities of M83 vs K83 or T83 in human MxB? First, we considered the possibility that M83 could function as an alternative translational start site, yielding an N-terminally truncated protein that is uniquely capable of inhibiting HSV-1. Previous studies have reported one such downstream initiation codon at methionine 26 in human MxB, which produces a short isoform of MxB [14]. However, this short form does not inhibit HIV-1 or HSV-1, suggesting the N-terminus is necessary for the antiviral function of MxB, possibly because it is needed for correct localization [7, 10, 22]. Our studies also did not reveal a clear band corresponding to the expected size of a protein produced by translational initiation at codon 83 on our immunoblots, although we cannot rule out the possibility that even at low abundance, such an isoform could correctly localize, either by itself or via interaction with full length MxB, to form a potent inhibitor of HSV-1. Future studies will help reveal whether methionine at codon 83 is unique in its ability to confer HSV-1 restriction, or if other amino acid substitutions might also retain antiviral activity, providing more clarity into this possibility.

Second, we considered the possibility of differential protein stability of MxB either due to inherent protein stability or post-translational modifications at residue 83 [23]. The HSV-1 immediate early ICP0 protein is known to facilitate the degradation of PML and ND10 via its E3 ligase activity [24]. K83, which is found in almost all primate MxBs, could similarly make MxB a target for ubiquitination and lead to its degradation during HSV-1 infection. However, when we analyzed human and chimp MxB expression in uninfected vs infected cells, we found that the levels of both orthologs remained similar regardless of infection status (Fig. 5), arguing against the differential degradation possibility.

A third possibility is that residue 83 may lie directly at the interface between MxB and as yet unidentified target proteins encoded by HSV-1; M83 could enhance this interaction explaining the increased restriction. Unfortunately, the crystal structure of MxB was solved without the N-terminal 83 amino acids, which are predicted to be unstructured [15]. Nevertheless, our finding that codon 83 is critical for counteracting HSV-1 will facilitate further studies aimed at understanding the mechanisms by which human MxB blocks HSV-1 in particular, and herpesviral replication in general. Studies of HIV-1 have suggested that MxB interacts with incoming viral capsids to block entry into the nucleus [7]. Consistent with this model, certain mutations of the HIV-1 capsid protein conferred resistance to MxB [8]. Another recent paper using partially purified HSV-1 capsids and MxB expressing cell extracts revealed that MxB can ‘punch holes’ in herpesviral capsids, leading to the premature release of viral DNA into the cytoplasm, which could in turn activate DNA sensors capable of amplifying the antiviral immune response [25]. Interestingly, HSV-1 capsid-associated tegument proteins appear able to reduce the capsid damage caused by MxB. Since human MxB, but not orthologs from other primates, is able to antagonize HSV-1, it may be that human MxB has a unique ability to bind to the HSV-1 capsid via improved binding conferred by the M83 variant, or an improved capacity to circumvent the putative tegument protein shield [25].

Interestingly, codon 83 also represents the most polymorphic non-synonymous polymorphism in human MxB. Approximately 5% of over 280,000 sequenced MxB human alleles have a M83T polymorphism, whose frequency rises to ∼18% in populations with African ancestry (data from gnomeAD [19];[4]), with a significant proportion of individuals homozygous for the T83 variant. Although this geographical distribution is intriguing, previous F_st_ analysis (measure of population genetic differentiation) of human MxB alleles did not find evidence of selection-driven differentiation at this site, rs56680307 [4]. However, our demonstration that the major M83 variant can restrict HSV-1 whereas the minor T83 variant cannot, does carry strong implications for an increased natural susceptibility to HSV-1 replication in a significant subset of the human population. Moreover, it still leaves unclear what evolutionary pressure may have led to the apparent loss of the protective MxB allele in human populations. Nevertheless, our findings suggest that the majority of human hosts unusually appear to be in the ‘winning position’ in their arms race with HSV-1 by virtue of the acquisition of the M83 MxB variant in the common ancestor of humans. To fully understand the impact of this allele, it will be important to translate these insights of MxB restriction of HSV-1 in cell culture systems to its replication and spread in nature.

## Materials and Methods

### Cells

IMR90, Vero, HF, and U2OS cells were maintained in Dulbecco’s Modified Eagle’s medium supplemented with 10% NuSerum (BD Biosciences). U2OS osteosarcoma cells containing a loxP landing pad were obtained from Daphne Avgousti (Fred Hutchinson Cancer Center) [12]. IMR90 MxB KO cells were constructed by transducing the cells with lentiviral vector (lentiCRISPR v2; Addgene #52961) containing a gRNA targeting the genomic MxB target sequence (5’-GGTGGTGGTTCCCTGTAACG-3’) as previously described [26]. Following selection with puromycin (1 ug/mL), these cells were transduced with lentiviral vectors derived from pEQ1607 [27] and containing various MxB transgenes with silent mutations in the gRNA target region (described below). Ortholog-expressing cells were then selected with hygromycin B (100 ug/mL). U2OS cells expressing MxB orthologs under the control of a dox-inducible promoter were constructed by transfecting plasmid pDA0173 containing the different MxB orthologs along with Cre recombinase, and selecting with puromycin (1 ug/mL).

### Viruses, infections, and plaque assays

HCMV (strain AD169; ATCC VR-538) was propagated in HFs. HSV-1 strains KOS and 17 were obtained from Daphne Avgousti (Fred Hutchinson Cancer Center). HSV-2 strain MS was obtained from ATCC (VR-540). Both HSV-1 and HSV2 were propagated and titered on Vero cells. For MxB restriction assays cells were either pretreated with dox for 24h (1 ug/mL) or left untreated, and then infected with the indicated viruses at an MOI of 0.1 for 48h for HSV-1 and HSV2, and 6 days for HCMV. Supernatants were then collected, and viral replication was quantified by plaque assay. HSV-1 and HSV-2 plaque assays were performed on Vero cells by incubating dilutions of each sample for 48 hours and then staining with crystal violet. HCMV titers were determined on HFs by incubating dilutions of each sample for 6 days prior to staining with crystal violet.

### Plasmids

MxB orthologs from human, chimpanzee, African green monkey, and owl monkey, that containing a C-terminal 3x flag tag were previously cloned into the pQCXIP vector [6]. Using these plasmids as templates, we introduced silent mutations into each ortholog at the guide RNA target site using primers #2607 and #2608 in stitch PCR reactions. The outside primers were #2534 and #2543 for human and chimp MxB, #2536 and #2543 for AGM MxB, #2561 and #2543 for owl monkey MxB. The resulting amplicons were cloned into lentiviral vector pEQ1607 digested with BstEII [27] by Gibson assembly. The resulting constructs, pEQ1779 (human), pEQ1819 (AGM), pEQ1823 (owl monkey), and pEQ1817 (chimp) express MxB under the control of a dox-inducible promoter. To generate U2OS cells stably expressing the MxB orthologs, we PCR-amplified the orthologs contained in the pEQ1607-based vectors using primers #2654 and #2655 (human and chimp), #2653 and #2655 (AGM), and #2652 and #2655 (owl monkey). The products were then introduced into HiLo vector (DA0173 [12], provided by Daphne Avgousti) digested with Ippo-1 and BstX1 by Gibson assembly. The resulting constructs were pEQ1774 (human MxB), pEQ1782 (chimp MxB), pEQ1780 (AGM MxB), and pEQ1778 (owl monkey MxB), respectively. Plasmid sequences were verified by long-read sequencing (Plasmidsaurus, Eugene, Oregon). The resulting sequences revealed that some of the ortholog sequences originally deposited in GENBANK were missing 42 nt from the 3’ end of MxB. In addition, the chimp MxB ortholog was found to have two point mutations, one at codon 48 that resulted in an asparagine to isoleucine substitution not present in other chimp sequences (great ape genome project [28]).

MxB chimeras were generated by restriction digestion of the human and chimp MxB constructs (pEQ1774 and pEQ1782) using combinations of restriction enzymes BsrG1, Sph1, and Not1 (NEB). These fragments were then swapped into each parental vector using T4 DNA ligase (Invitrogen). Chimeras A and B (pEQ1785 and pEQ1784) were generated by cutting pEQ1774 and pEQ1782 with BsrG1 and Not1 and swapping the N-terminal segments. Chimeras C and D (pEQ1787 and pEQ1786) were generated by cutting pEQ1774 and pEQ1782 with BsrG1 and Sph1 and swapping the internal segments. Chimeras E and F (pEQ1791 and pEQ1788) were generated by cutting pEQ1774 and pEQ1782 with Sph1 and Not1 and swapping the C-terminal segments.

Point mutants were generated by PCR amplification of either pEQ1774 or pEQ1782 using additional internal primers to introduce the mutations. The resulting amplicons were then joined by overlap extension PCR. Chimp MxB I48N (pEQ1794) was generated by amplifying pEQ1782 with primers #2654 and #2882 (5’ end) and primers #2883 and #2655 (3’ end). Chimp MxB K83M (pEQ1795), was generated by amplifying pEQ1782 with primers #2654 and #2886 (5’ end) and primers #2887 and #2655 (3’ end). Human MxB M83K (pEQ1793), was generated by amplifying pEQ1774 with primers #2654 and #2888 (5’ end) and primers #2889 and #2655 (3’ end). Human MxB M83T (pEQ1804), was generated by amplifying pEQ1774 with primers #2654 and #2877 (5’ end) and primers #2878 and #2655 (3’ end). The amplicons were then Gibson cloned into the DAO173 HiLo vector and introduced into U2OS cells as described above. Table 1 contains a full list of the sequences of the primers used here.

### Immunoblot analyses

Samples for immunoblotting were prepared by washing cells with PBS, then lysing them in 2% SDS. The protein lysates were separated by 10% SDS-polyacrylamide gel electrophoresis on gels containing 0.5% 2,2,2-trichloroethanol to allow for stain free visualization of proteins [29], and then transferred to polyvinylidene difluoride (PVDF) membranes (Millipore). Blots were probed with the indicated antibodies using the Western Star chemiluminescence detection system (Applied Biosciences) according to the manufacturer’s recommendations. The antibodies used in these experiments include MxB (Abcam 196833), alkaline phosphatase-conjugated flag M2, flag, and actin (all from Sigma). Antibodies were used per the manufacturer’s recommendations. Immunoblot images were captured and quantified with a ChemiDoc Touch imaging system and Image Lab software (Bio-Rad Laboratories, Hercules, CA).

### Immunofluorescence

Cells were grown in 8 well chamber slides (LAB-TEK II) and either untreated or treated with doxycycline for 24h. Cells were then fixed with 0.4% paraformaldehyde, and permeabilized with 0.25% triton x-100. The slides were incubated in blocking agent for 1hr, with flag antibody (1:10,000) for 1h, rinsed, then incubated with an anti-mouse 555 fluorophore (Santa Cruz). Images were captured on a high-resolution Leica Stellaris Confocal Microscope using a 63x oil objective and analyzed using ImageJ software.

### Statistical Analysis

All experiments were repeated a minimum of 3 times. Statistical analyses were performed using a Student’s T test to calculate statistical significance. ns P>0.05; * = P< 0.05; ** = P< 0.01; *** = P<0.001; **** = P<0.0001. Statistical analyses were performed using Prism 7 software (GraphPad).

## Acknowledgements

We thank Daphne Avgousti and Michael Emerman (both at Fred Hutchinson Cancer Center) for reagents and helpful comments. We also thank the Fred Hutchinson Cancer Center Genomics Shared Resource for technical assistance. This work was supported by NIH RO1 AI145945 (to APG), NIH T32 CA080416 (to AB), Howard Hughes Medical Institute (HHMI, to HSM), and by the Fred Hutch/University of Washington Cancer Consortium (P30 CA015704). HSM is an investigator of HHMI. The content is solely the responsibility of the authors and does not necessarily represent the official views of the National Institutes of Health. The funders had no role in study design, data collection and analysis, or preparation of the manuscript.

---

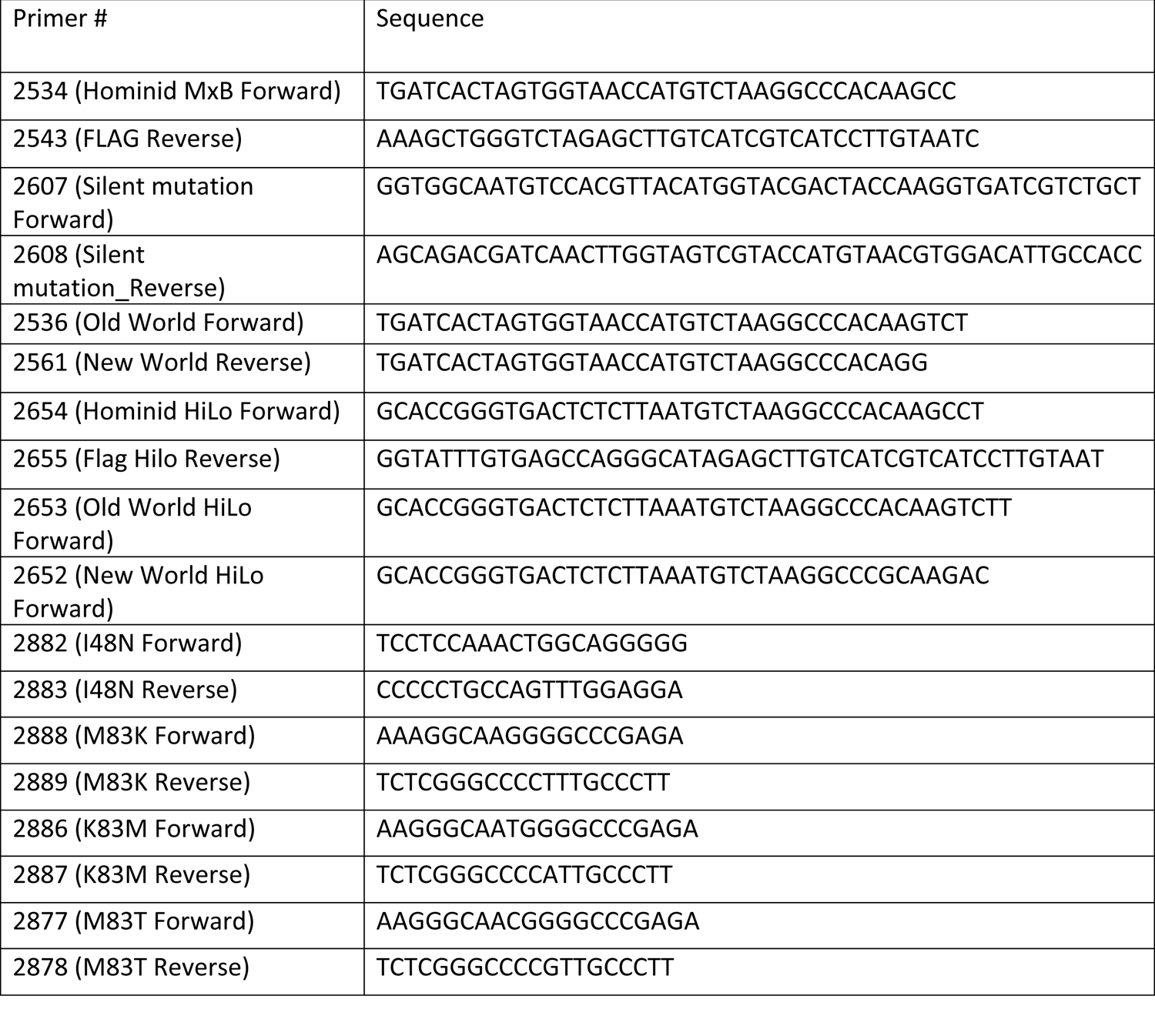

## Notes

### Competing Interest Statement

The authors have declared no competing interest.

